# Isolation of SAR11 marine bacteria from cryopreserved seawater

**DOI:** 10.1101/2020.09.22.309336

**Authors:** Elizabeth A. Monaghan, Kelle C. Freel, Michael S. Rappé

## Abstract

In this study, we sought a means to increase current culture collections of SAR11 marine bacteria by testing the use of seawater cryopreserved with glycerol as an inoculum. In July 2017, raw seawater was collected outside of Kāne‘ohe Bay, Hawai‘i, in the tropical Pacific Ocean. A portion of this sample was diluted in seawater-based growth medium to create 576 × 2 mL dilution cultures containing 5 cells each and incubated for a high-throughput cultivation experiment, while another portion was cryopreserved in 10% glycerol. After ten months, a cryopreserved aliquot of seawater was thawed, diluted in seawater-based growth medium, and distributed to create a second high-throughput cultivation experiment of 480 × 2 mL dilution cultures containing 5 cells each and 94 cultures containing 105 cells each. The raw seawater cultivation experiment resulted in the successful isolation of 54 monocultures and 29 mixed-cultures, while cryopreserved seawater resulted in 59 monocultures and 29 mixed cultures. Combined, the cultures included 51 SAR11 isolates spanning 11 unique 16S rRNA gene amplicon sequence variants (ASVs) from raw seawater inoculum and 74 SAR11 isolates spanning 13 unique ASVs from cryopreserved seawater. A vast majority (115 of 125) of SAR11 isolates from the two HTC experiments were members of SAR11 subclade Ia, though isolates of subclades IIIa and Va were also recovered from cryopreserved seawater and subclade Ib was recovered from both. The four most abundant SAR11 subclade Ia ASVs found in the initial seawater sample used to create both culture experiments were isolated by both approaches.

**Importance:** High-throughput dilution culture has proved to be a successful approach to bring some difficult-to-isolate planktonic microorganisms into culture, including the highly abundant SAR11 lineage of marine bacteria. While the long-term preservation of bacterial isolates by freezing in the presence of cryoprotectants such as glycerol has been shown to be an effective method of storing viable cells over long time periods (i.e. years), to our knowledge it had not previously been tested for its efficacy in preserving raw seawater for later use as inoculum for high-throughput cultivation experiments. We found that SAR11 and other abundant marine bacteria could be isolated from seawater that was previously cryopreserved for nearly 10 months, at a rate of culturability similar to that of the same seawater used fresh, immediately after collection. Our findings expand the potential of high-throughput cultivation experiments to include opportunities where immediate isolation experiments are impractical, allow for targeted isolation experiments from specific samples based on analyses such as microbial community structure, and enable cultivation experiments across a wide range of other conditions that would benefit from having source inoculum available over extended periods of time.

## Introduction

The rapid advancement of molecular tools to investigate marine microorganisms in their natural environment has led to unprecedented access to the genomic repertoire and transcription- and protein-based assessments of activity within natural microbial cells, populations, and communities (1, 2). It is currently feasible for a few liters of seawater to provide sequence data that reveals microbial population structure, the identities of microbial community members, and the presence and activity of potential metabolic functions they harbor (e.g. 3, 4). In recent years, however, the value of having cultivated representatives of numerically abundant and environmentally relevant microbial lineages has received renewed recognition (5–8). Access to isolated strains or low-diversity enrichments of marine microorganisms that are commonly found in the natural environment has provided a means to definitively test many hypotheses generated from environmental observations and experiments, as well as whole genome sequences useful for informing and guiding environmental genomics, transcriptomics, and proteomics research (e.g. 5, 6). The importance of cultivating environmentally-relevant microorganisms from pelagic marine ecosystems for laboratory-based experimentation is now generally appreciated. However, evidence provided by the sequencing of environmental DNA continues to support the conclusion that most of the microorganisms that appear to dominate pelagic marine ecosystems have not yet been cultivated from seawater (11).

Several different isolation methods and strategies have been developed in order to coax recalcitrant environmental microorganisms into laboratory culture (e.g 12–15). Among these novel approaches is an isolation technique based on dilution-to-extinction culturing methodology first developed by Button and colleagues (16). Although early dilution-to-extinction culturing studies resulted in cultures of novel oligotrophs (16–18), the dilution culture strategy was not without limitations. For instance, the technique yielded only a small number of isolates, while requiring a significant amount of time and effort per experiment. The high-throughput culturing (HTC) approach is a variation of dilution-to-extinction culturing methodology tailored to facilitate rapid, high-throughput experiments with high rates of replication and greater opportunities to investigate physical, chemical, and biological variables (19, 20).

For over a decade since its initial discovery in 1990, the marine planktonic bacterial lineage known as SAR11 served as a notorious example of an abundant and widespread microorganism in nature that was recalcitrant to cultivation as an isolated strain in a controlled laboratory setting (21, 22). The value of the HTC strategy was solidified when early trials yielded the first cultured representatives of many marine microbial groups that were previously known only from environmental SSU rRNA genes (19), including the first cultivated strains of SAR11 (20). In general, the HTC approach employs growth media created from natural or artificial seawater in order to dilute the cells within a fluid sample, which is then arrayed in high density replicate cultures, propagated, and monitored under controlled conditions. In addition to diverse SAR11 strains (20, 23–28), the application of this method has resulted in the isolation of numerous other important lineages of marine bacteria including OM43 (19, 29), SAR116 (23, 30), SAR92 (23, 31), and SUP05 (32), among others. While efforts have succeeded in isolating many abundant planktonic marine bacteria, it remains that the genetic diversity harbored by these lineages in nature greatly surpasses what has been isolated in the laboratory.

A current limitation of the HTC approach is that, thus far, it has only been used with freshly collected inoculum. This presents a potential constraint on HTC experiments using fluid samples collected in the field as it requires all of the resources necessary for setting up an HTC experiment (appropriate laboratory space, biosafety cabinet, etc.) be available at or near the time and location of sampling. The preservation of cultivated bacterial strains by freezing in the presence of cryoprotectants such as glycerol or dimethyl sulfoxide (DMSO) has proved an effective method for preserving viable cells over long time periods (i.e. years), including cultivated strains of SAR11 (20, 33, 34). However, to our knowledge it has not previously been tested for its efficacy in preserving raw seawater for subsequent use as inoculum for high-throughput cultivation. In this study, we conducted HTC experiments to compare the use of a raw seawater sample collected from the coast of O‘ahu, Hawai’i, in the tropical Pacific Ocean, with a subsample of the same seawater that was cryogenically preserved for nearly ten months. In particular, we sought to determine if members of the SAR11 lineage of marine bacteria could be isolated from cryopreserved seawater and thus open the possibility to expand existing culture collections of SAR11 to potentially include any locations where seawater samples could be collected and preserved.

## Results

### Overview of HTC cultivation experiments

Using seawater sampled outside of Kāne‘ohe Bay on the island of O‘ahu, Hawai‘i (Fig. 1), two high-throughput cultivation experiments were conducted: one that used fresh seawater as an inoculum, labeled HTC2017, and one that used a cryopreserved sample of the same seawater ∼10 months later (HTC2018) (Fig. 2). Of 576 initial 2 mL cultures inoculated with raw seawater for the HTC2017 experiment, 150 exhibited positive growth after 56 days of incubation. Of these, 123 contained sufficient volume of culture to be sub-cultured into 20 mL of fresh medium. Following DNA extraction, sequencing, and the assignment of ASVs, 54 monocultures and 29 mixed cultures were recovered (Table 1). The remainder either did not yield an amplification product or did not contain an ASV ≥50% of the culture and thus were not considered further. Fifty-four isolates were identified in the 29 mixed cultures (Table S1); the 108 unique isolates identified in the HTC2017 experiment (monocultures plus isolates contained in mixed cultures) were distributed amongst 28 ASVs in total (Table 2, Table S1). The HTC2017 experiment yielded a culturability of 3.1% (2.5% – 3.9%) when both monocultures and mixed cultures were considered, and 2.0% (1.5% – 2.6%) when considering only monocultures (Table 1).

**Table 1.**
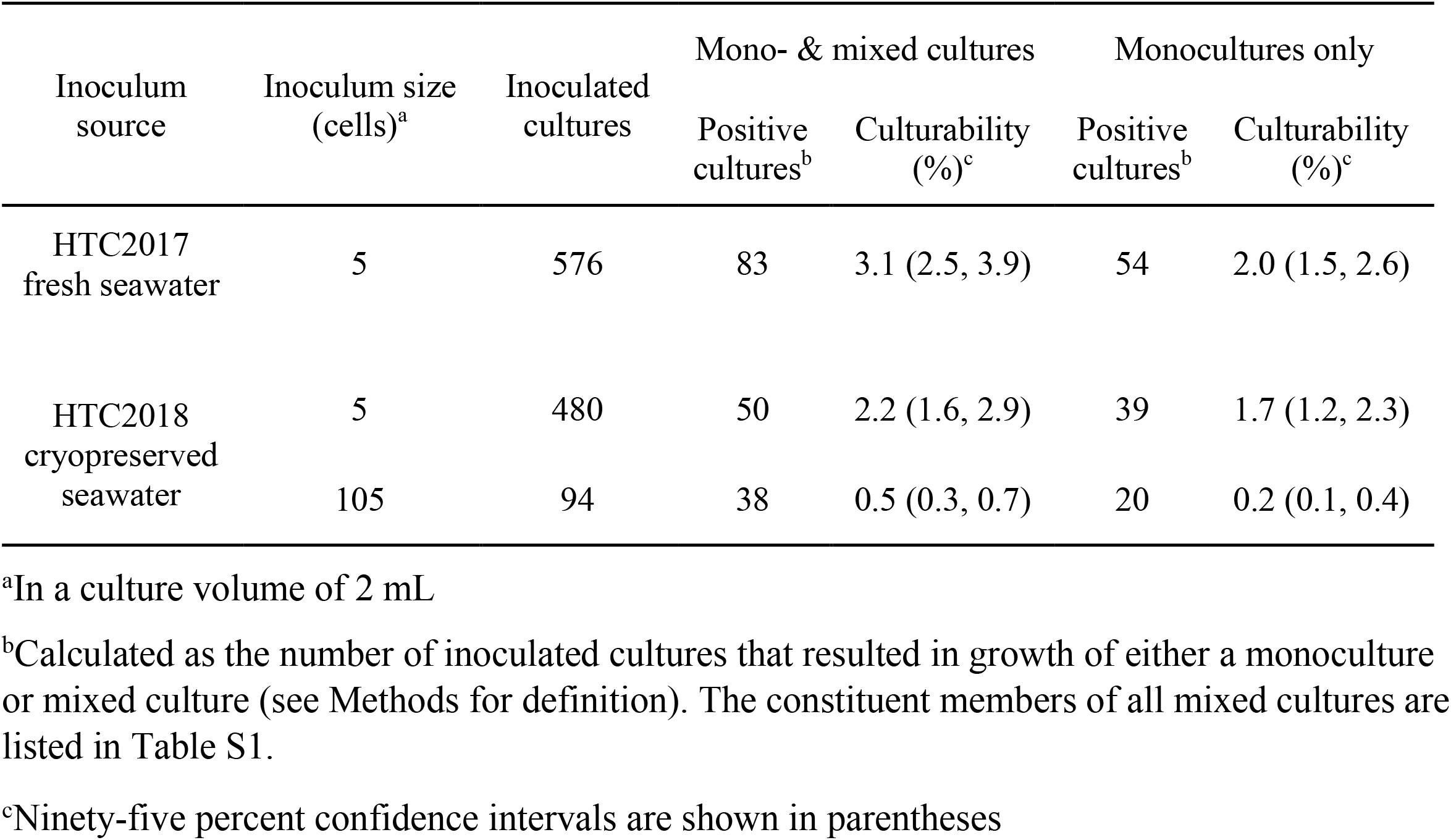
Culturability statistics from fresh (HTC2017) and cryopreserved (HTC2018) seawater cultivation experiments.

| Inoculum source | Inoculum size (cells)<sup>a</sup> | Inoculated cultures | Mono- & mixed cultures |  | Monocultures only |  |
| --- | --- | --- | --- | --- | --- | --- |
|  |  |  | Positive cultures<sup>b</sup> | Culturability (%)<sup>c</sup> | Positive cultures<sup>b</sup> | Culturability (%)<sup>c</sup> |
| HTC2017<br>fresh seawater | 5 | 576 | 83 | 3.1 (2.5, 3.9) | 54 | 2.0 (1.5, 2.6) |
| HTC2018<br>cryopreserved<br>seawater | 5 | 480 | 50 | 2.2 (1.6, 2.9) | 39 | 1.7 (1.2, 2.3) |
|  | 105 | 94 | 38 | 0.5 (0.3, 0.7) | 20 | 0.2 (0.1, 0.4) |

<sup>a</sup>In a culture volume of 2 mL
<sup>b</sup>Calculated as the number of inoculated cultures that resulted in growth of either a monoculture or mixed culture (see Methods for definition). The constituent members of all mixed cultures are listed in Table S1.
<sup>c</sup>Ninety-five percent confidence intervals are shown in parentheses

**Figure 1.**
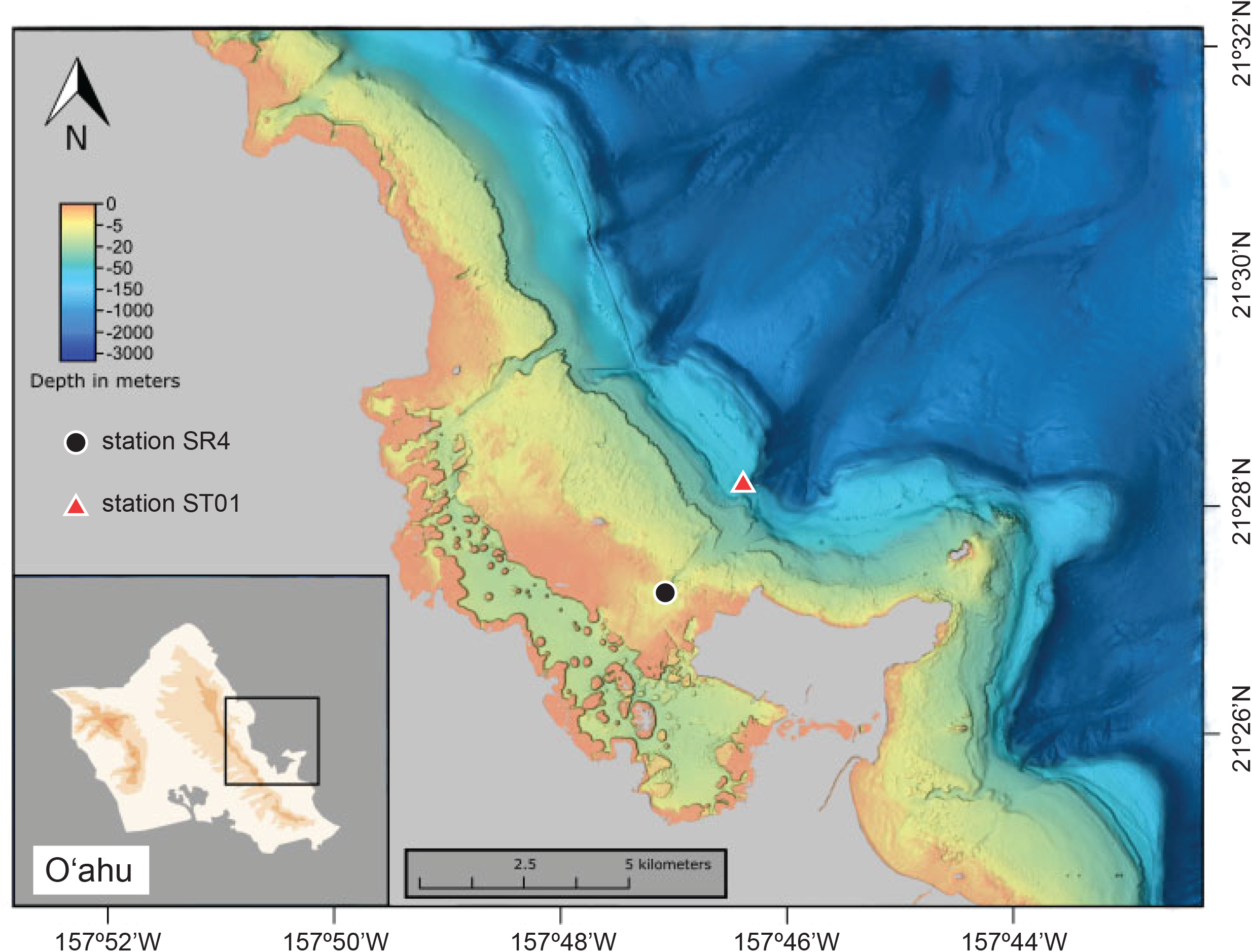
Map indicating the location of stations STO1 (red triangle) and SR4 (black circle) in the vicinity of Kāne‘ohe Bay on the island of O‘ahu, Hawai‘i, where seawater used as inoculum (STO1) and media preparation (SR4) were collected.

**Figure 2.**
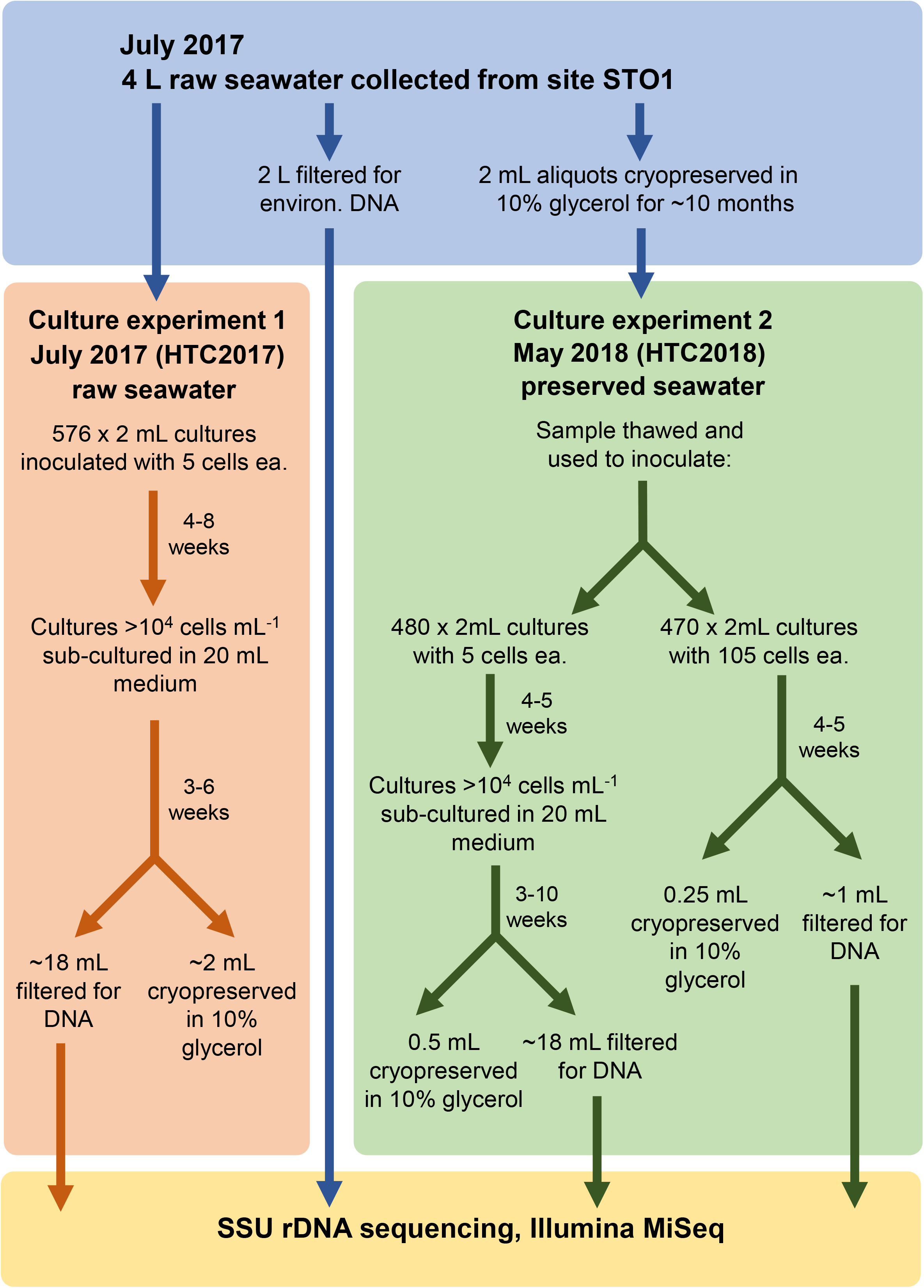
Outline of experiments performed in this study.

**Table 2.** Summary of isolates from fresh (HTC2017) and cryopreserved (HTC2018) seawater cultivation experiments, including the relative abundance of each taxonomic group in the environmental sample based on SSU rRNA gene sequencing.

| Taxonomy<sup>a</sup> | HTC2017 |  | HTC2018 |  | Seawater<sup>b</sup> |  |
| --- | --- | --- | --- | --- | --- | --- |
|  | ASVs | Strains | ASVs | Strains | ASVs | Abundance (rel. %) |
| Alphaproteobacteria, SAR11 subclade Ia | 9 | 47 | 9 | 68 | 10 | 10.23 |
| Alphaproteobacteria, SAR11 subclade Ib | 2 | 4 | 2 | 3 | 14 | 7.74 |
| Alphaproteobacteria, SAR11 subclade IIIa | 0 | 0 | 1 | 2 | 4 | 0.42 |
| Alphaproteobacteria, SAR11 subclade Va | 0 | 0 | 1 | 1 | 2 | 2.13 |
| Alphaproteobacteria, Rhodobacteraceae | 4 | 14 | 5 | 5 | 12 | 5.81 |
| Alphaproteobacteria, SAR116 clade | 1 | 1 | 0 | 0 | 23 | 5.41 |
| Alphaproteobacteria, PS1 clade | 1 | 3 | 3 | 4 | 3 | 0.42 |
| Gammaproteobacteria, Halieaceae, OM60(NOR5) clade | 4 | 24 | 5 | 21 | 5 | 1.91 |
| Gammaproteobacteria, KI89A clade | 1 | 1 | 0 | 0 | 4 | 0.60 |
| Betaproteobacteriales, Burkholderia-Caballeronia-Paraburkholderia | 1 | 4 | 1 | 1 | 2 | 0.11 |
| Betaproteobacteriales, Methylophilaceae, OM43 clade | 2 | 5 | 0 | 0 | 1 | 0.04 |
| Gammaproteobacteria, Pseudomonadaceae, Pseudomonas | 1 | 2 | 0 | 0 | 0 | 0 |
| Gammaproteobacteria, Rhodanobacteraceae | 1 | 2 | 0 | 0 | 0 | 0 |
| Actinobacteria, Corynebacteriaceae | 0 | 0 | 1 | 1 | 0 | 0 |
| Actinobacteria, Geodermatophilaceae | 0 | 0 | 1 | 1 | 0 | 0 |
| Bacteroidetes, Chitinophagaceae, Sediminibacterium | 0 | 0 | 1 | 1 | 0 | 0 |
| Fungi | 1 | 1 | 1 | 2 | 0 | 0 |

<sup>a</sup>Abbreviated taxonomy adapted from Silva release 132; the full taxonomy of each ASV is available in Table S1.
<sup>b</sup>Environmental relative abundance was calculated using read counts of all environmental ASVs detected in the seawater sample from within each taxonomic grouping, after curation to remove ASVs originating from chloroplasts.

For the cryopreserved seawater experiment (HTC2018), wells were inoculated with either 5 or 105 cells. Of the 480 initial 2 mL cultures inoculated with 5 cells well^−1^ of cryopreserved seawater (HTC2018), 142 exhibited positive growth after 30 days of incubation. The 142 positive wells were subcultured into 20 mL seawater media and, after 72 days, 95 subcultures ultimately exhibited growth. Following DNA extraction, sequencing, and the assignment of ASVs, 39 monocultures and 11 mixed cultures were identified (Table 1). The remainder either did not yield an amplification product or did not contain an ASV ≥ 50% of the culture and thus were not considered further. Sixteen isolates were identified in the 11 mixed cultures (Table S1). Combined, the 55 unique isolates identified in the HTC2018 5 cells well^−1^ experiment were distributed amongst 17 ASVs (Table 2, Table S1). The HTC2018 5 cells well^−1^ experiment yielded a culturability of 2.2% (1.7% – 2.9%) when both monocultures and mixed cultures were considered, and 1.7% (1.2% – 2.3%) when considering only monocultures (Table 1).

Of the 470 initial 2 mL cultures inoculated with 105 cells well^−1^ of cryopreserved seawater (HTC2018), 343 exhibited positive growth after 31 days of incubation. A single 96-well cultivation plate containing 64 positive wells and two uninoculated control wells were selected for further processing. Following DNA extraction, sequencing, and the assignment of ASVs, 20 monocultures and 18 mixed cultures were identified (Table 1). The remainder either did not yield an amplification product or did not contain an ASV ≥ 50% of the culture and thus were not considered further. Thirty-five isolates were identified in the 18 mixed cultures (Table S1); combined, the 55 unique isolates identified in the 105 cells well^−1^ HTC2018 experiment were distributed amongst 21 ASVs (Table 2, Table S1). The HTC2018 105 cells well^−1^ experiment yielded a culturability of 0.5% (0.3% – 0.7%) when both monocultures and mixed cultures were considered, and 0.2% (0.1% – 0.4%) when considering only monocultures (Table 1).

### Identity of isolates

After quality control, each culture was sequenced to an average depth of 14,047 ± 8,014 (s.d.; range of 679 – 57,557) reads. Regardless of whether they originated from raw or cryopreserved seawater, the broad, bacterial family-level taxonomic identity of isolates revealed substantial overlap between culture experiments (Table 2, Table S1). Members of the alphaproteobacterial SAR11 subclade I, the marine gammaproteobacterial family *Halieaceae*, and the alphaproteobacterial family *Rhodobacteraceae* were the first-, second-, and third-most abundant families isolated in both the fresh seawater (HTC2017) and cryopreserved seawater (HTC2018) cultivation experiments (Table 2, Table S1). Combined, these three groups made up 82% (89 of 108) and 88% (97 of 110) of isolates recovered from HTC2017 and HTC2018, respectively. Other bacterial families with isolates shared between HTC2017 and HTC2018 include the PS1 clade of *Alphaproteobacteria* and *Burkholderiaceae* within the *Betaproteobacteria* (Table 2, Table S1). When the five bacterial families shared between HTC2017 and HTC2018 are considered, the fresh seawater and cryopreserved seawater cultivation experiments shared 89% (96 of 108) and 93% (102 of 110) of isolated strains, respectively.

#### SAR11

A total of 51 strains representing 11 unique ASVs of SAR11 marine bacteria (alphaproteobacterial order *Pelagibacterales*) were cultivated in HTC2017, while 74 strains representing 13 ASVs were cultivated in HTC2018 (Fig. 3, Table 2, Table S1). They make up 47% and 67% of the isolates recovered in the two experiments, respectively. The vast majority of these isolates were members of SAR11 subclade Ia, including 47 strains from HTC2017 and 68 strains from HTC2018. Each experiment resulted in the isolation of nine subclade Ia ASVs, including five that were common between the two experiments (Figs. 3 & 4, Table S1). The two most often isolated SAR11 subclade Ia ASVs were shared between the two experiments: ASV003 (25 and 37 isolates) and ASV002 (11 and 13 isolates) from HTC2017 and HTC2018, respectively (Fig. 3, Table S1). Two other subclade Ia ASVs (ASV034 and ASV046) consisted of multiple strains from both experiments, while 7 subclade Ia ASVs consisted of a single isolate from one experiment (Fig. 3, Table S1).

**Figure 3.**
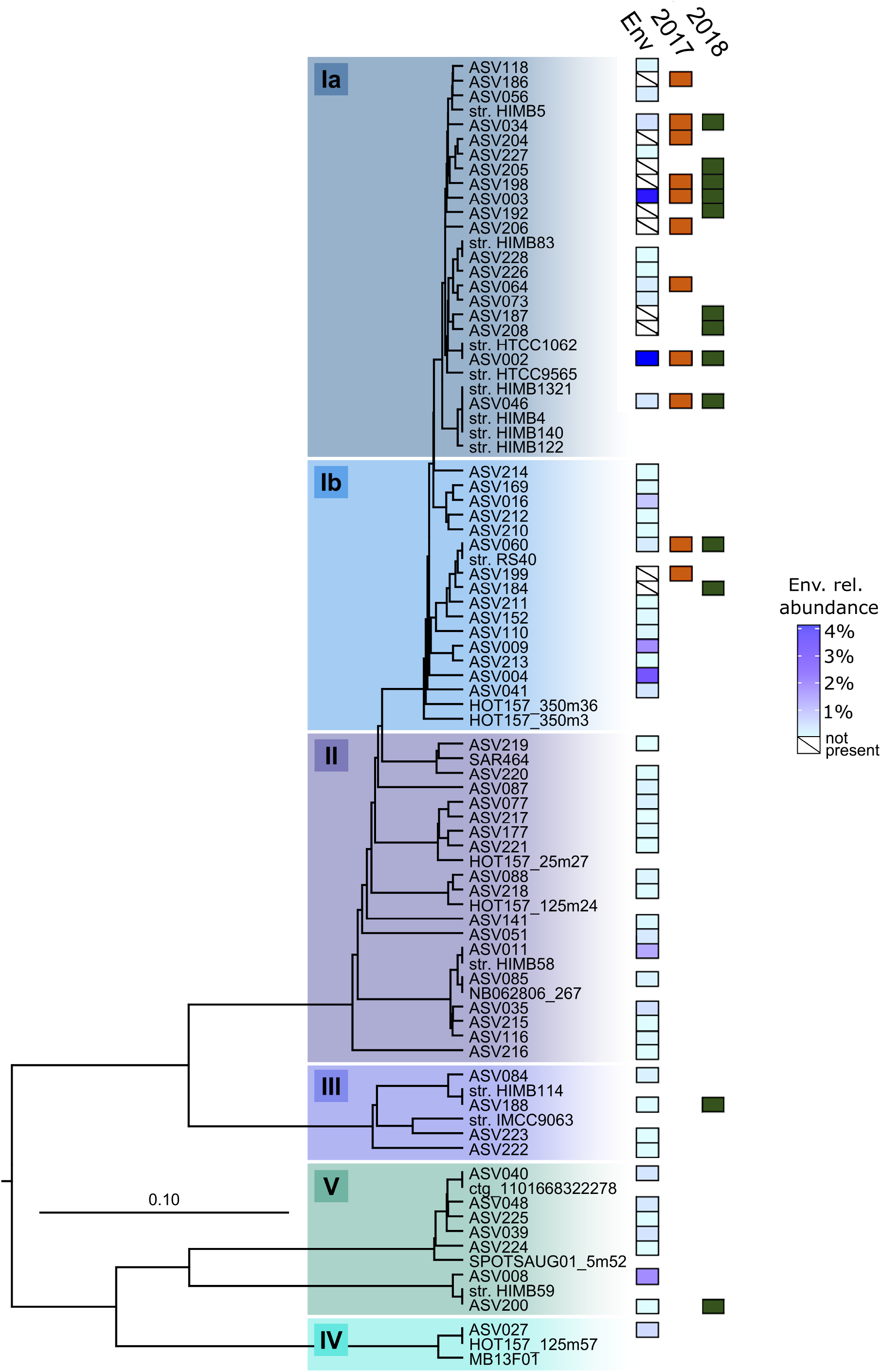
Phylogenetic analysis of the SAR11 clade illustrating relationships among 16S rRNA gene ASVs recovered from isolates and the source seawater used as inoculum in this study. The scale bar corresponds to 0.1 substitutions per nucleotide position. A variety of *Alphaproteobacteria* were used as an outgroup. Previously cultured isolates (“str.”) and select environmental gene clones were included as references. Boxes labeled “Env” indicate the relative environmental abundance of each ASV (blue gradient), while orange (“2017”) and green (“2018”) boxes indicate the presence of an ASV in the fresh (HTC2017) and cryopreserved (HTC2018) seawater cultivation experiments respectively. Boxes containing a slash in the “Env” column indicate ASVs was found in a culture but were not detected in the environmental sample.

Strains affiliated with SAR11 subclade Ib were also isolated from both fresh and cryopreserved seawater, including 4 isolates across 2 ASVs from HTC2017 and 3 isolates across 2 ASVs from HTC2018 (Fig. 4, Table S1). SAR11 subclade Ib ASV060 consisted of 3 and 2 isolates from HTC2017 and HTC2018, respectively, while each experiment also yielded an isolate with a unique subclade Ib ASV (Fig. 3, Table S1). Two SAR11 subclades were only isolated from the cryopreserved seawater sample, including two isolates from subclade IIIa (ASV188) and one isolated from subclade Va (ASV200) (Fig. 3, Table 2, Table S1).

**Figure 4.**
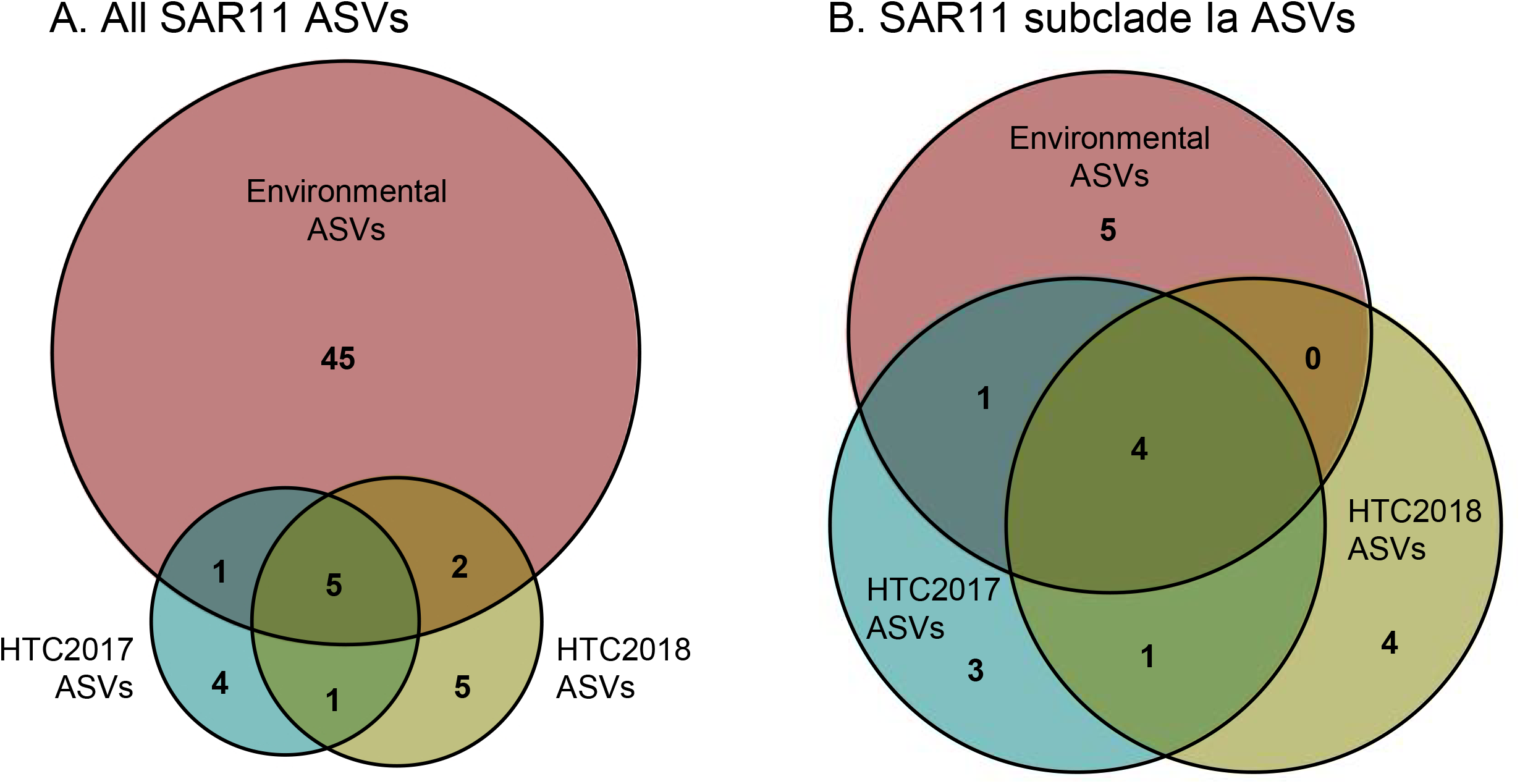
Venn diagrams comparing (A) SAR11 ASVs identified within the environmental seawater sample used as inoculum, isolates from the fresh seawater cultivation experiment (HTC2017), and isolates from the cryopreserved seawater cultivation experiment (HTC2018). (B) Same as (A), except limited to SAR11 subclade Ia ASVs.

#### OM60(NOR5)

Within the gammaproteobacterial family *Halieaceae*, the marine OM60(NOR5) clade made up 24 and 21 isolates, or 22% and 19% of HTC2017 and HTC2018, respectively (Table 2). The 4 ASVs that accounted for the 24 isolates from HTC2017 were shared with HTC2018, where they accounted for 19 of the 21 isolates recovered from that experiment (Fig. 5, Table S1). One additional OM60(NOR5) ASV (ASV201) consisting of 2 isolates was recovered from cryopreserved seawater (Fig. 5, Table S1). Two closely related ASVs (ASV032 and ASV018) accounted for most of the OM60(NOR5) strains isolated from both experiments of this study (Fig. 5, Table S1). ASV32 was identical to strain HIMB55, a genome-sequenced member of the OM60(NOR5) clade previously isolated from Kāne‘ohe Bay, Hawai‘i (35). ***Rhodobacteraceae***. The marine alphaproteobacterial family *Rhodobacteraceae* made up 14 and 5 isolates, or 13% and 5% of HTC2017 and HTC2018, respectively (Table 2). The strains were distributed amongst 4 (HTC2017) and 5 (HTC2018) ASVs, including three (ASV12, ASV71, ASV124) that were shared between the two experiments (Fig. 5, Table S1). Eight of 14 *Rhodobacteraceae* isolates recovered from HTC2107 belonged to a single ASV (ASV012). ASV71 was identical to strain HIMB11, a genome-sequenced member of the *Rhodobacteraceae* previously isolated from Kāne‘ohe Bay, Hawai‘i (Fig. 5) (36).

**Figure 5.**
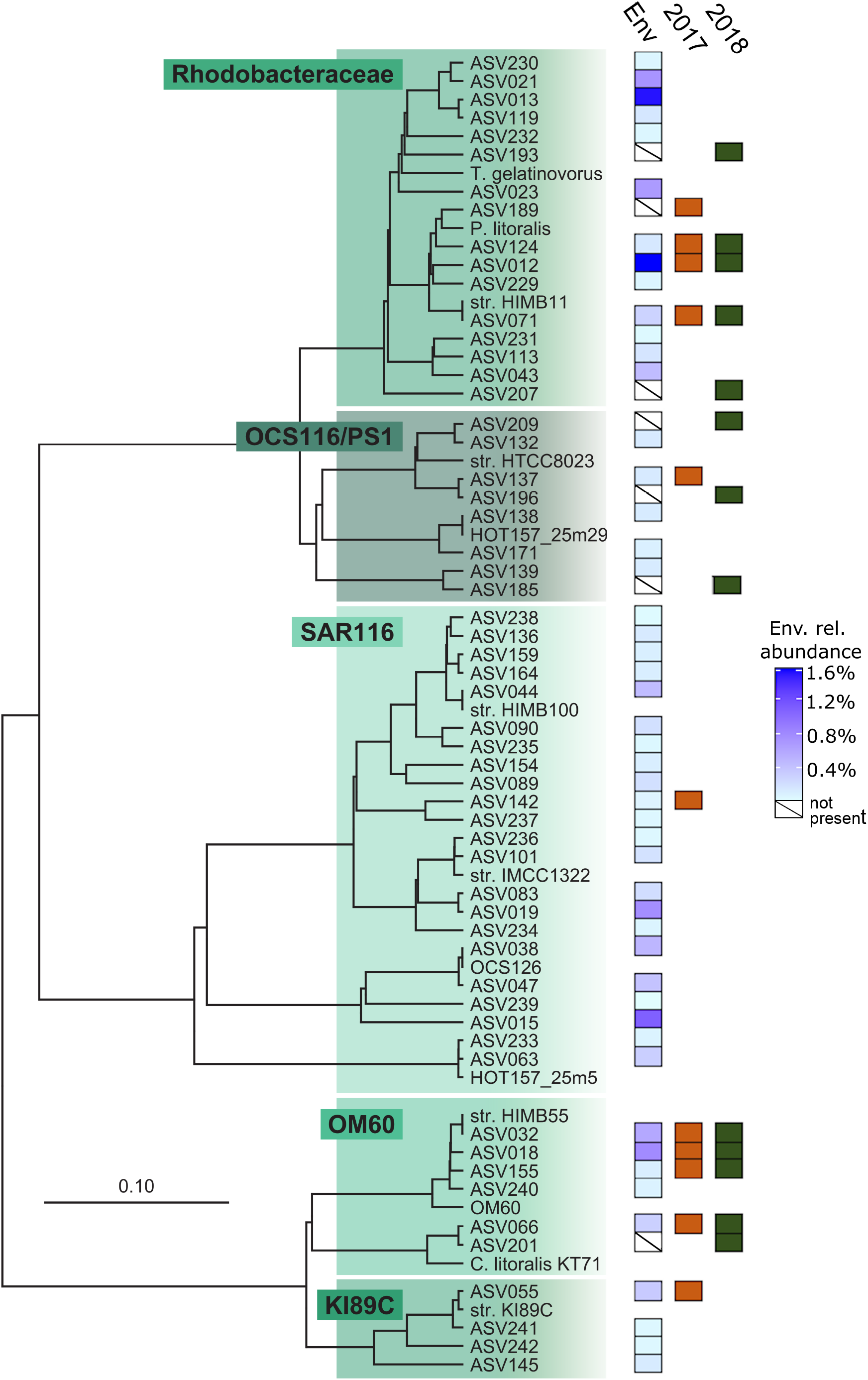
Phylogenetic analysis of select lineages of *Alpha*- and *Gammaproteobacteria* illustrating relationships among 16S rRNA gene ASVs recovered from isolates and the source seawater used as inoculum in this study. The scale bar corresponds to 0.1 substitutions per nucleotide position. A variety of *Betaproteobacteria* were used as an outgroup. Previously cultured isolates (“str.”) and select environmental gene clones were included as references. Boxes labeled “Env” indicate the relative environmental abundance of each ASV (blue gradient), while orange (“2017”) and green (“2018”) boxes indicate the presence of an ASV in the fresh (HTC2017) and cryopreserved (HTC2018) seawater cultivation experiments respectively. Boxes containing a slash in the “Env” column indicate ASVs was found in a culture but were not detected in the environmental sample.

#### Other isolates

In addition to the SAR11, OM60(NOR5), and *Rhodobacteraceae* ASVs described above, one additional ASV was isolated in both experiments: ASV190 within the betaproteobacterial family *Burkholderiaceae* was represented by 4 strains in HTC2017 and 1 in HTC2018 (Table S1). While identical ASVs were not isolated, members of the marine alphaproteobacterial PS1 clade within the order *Parvibaculales* were recovered from both cultivation experiments, including 3 isolates from a single ASV (ASV137) in HTC2017 and 4 isolates from 3 ASVs in HTC2018 (Fig. 5, Table S1). The remaining ASVs were recovered as either singletons or pairs of strains except for the OM43 clade of the *Betaproteobacteriales*, which constituted five strains across 2 ASVs from the fresh seawater inoculum (HTC2017) only (Table S1).

### Comparisons with inoculum microbial community

After quality control, the 16S rRNA gene amplicon from the inoculum seawater sample was sequenced to a depth of 65,924 reads. This sample harbored a microbial community dominated by typical marine bacteria, including the marine picocyanobacteria *Prochlorococcus* and *Synechococcus*, multiple subclades of the SAR11 lineage, the family *Flavobacteriaceae* of the bacterial phylum *Bacteroidetes*, diverse members of the gammaproteobacterial SAR86 and OM60(NOR5) lineages and alphaproteobacterial SAR116 and *Rhodobacteraceae* lineages, and *Actinomarinaceae* of the bacterial phylum *Actinobacteria*, among others (Table S1).

Within the inoculum sample, 53 SAR11 ASVs totaling 26% of the microbial community were identified. These spanned a diverse array of subclades that included Ia, Ib, IIa, IIIa, IV, Va, and Vb (Fig. 3, Table S1). Eight of the 53 SAR11 ASVs were isolated from at least one cultivation experiment, including ASVs within subclades Ia, Ib, IIIa, and Va, while 5 of the 8 were isolated from both HTC2017 and HTC2018 (Figs. 3 & 4, Table S1). Of 10 SAR11 subclade Ia ASVs present in the inoculum, the four most abundant were cultivated from both experiments (ASV002, ASV003, ASV034, and ASV046), including the second- and third-most abundant individual ASVs in the inoculum seawater community (Table S1). A fifth was cultivated in HTC2017 only (Fig. 3, Table S1). Eight SAR11 subclade Ia ASVs were isolated that did not appear in the inoculum seawater community (Figs. 3 & 4, Table S1).

In contrast to SAR11 subclade Ia, other SAR11 subclades present in the inoculum seawater microbial community were cultivated rarely or not at all. For example, only 1 of 14 subclade Ib ASVs that appeared in the inoculum was isolated (ASV060; Fig. 3, Table S1), although it was cultivated in both HTC2017 and HTC2018 experiments. From the HTC2018 experiment, one of four subclade IIIa ASVs (ASV188) was isolated, as well as one of two subclade Va ASVs (ASV200; Fig. 3, Table S1). Thirteen *Rhodobacteraceae* ASVs were identified in the environmental sample, of which the same three (ASV012, ASV071, ASV124) were cultivated from both fresh and cryopreserved seawater (Fig. 5, Table S1). Four of the five total most environmentally abundant OM60(NOR5) clade ASVs were also cultivated in both experiments (Table S1). Despite 19 ASVs appearing in the inoculum, only one SAR116 ASV (ASV142; Fig. 5, Table S1) was cultivated in the HTC2017 experiment.

### Mixed cultures

Twenty-nine mixed cultures were identified within each of the HTC2017 and HTC2018 experiments, yielding a total of 58 mixed cultures (Table S1). A majority of the mixed cultures contained ASVs from either SAR11 subclade Ia or the OM60(NOR5) clade, which is logical given the high recovery of monocultures from these two groups in both cultivation experiments (Fig. S1). Of nine mixed cultures containing OM60(NOR5) ASV018, eight also contained SAR11 subclade Ia ASV002, ASV003, or ASV034 (Fig. S1, Table S1). Eight of the 11 mixed cultures containing OM60(NOR5) ASV032 also contained a SAR11 subclade Ia ASV as well (Fig. S1, Table S1). All cultivated OM43 clade ASVs were in mixed cultures; both of the OM43 clade ASV202 isolates appeared in co-culture with the OM43 clade ASV195 (Fig. S1, Table S1).

## Discussion

For a variety of reasons, SAR11 marine bacteria remain a target for culturing experiments, despite being first isolated nearly 20 years ago (20). In large part, this is driven by the enormous genomic diversity harbored by this lineage and the probability for ecotypic differentiation across the SAR11 phylogenetic tree (3, 26, 37, 38). Living cultures offer a direct means to characterize and quantify the cellular and physiological features that underly differences in abundance or activity observed via direct environmental sampling (4, 39, 40). The primary goal of this study was to test the hypothesis that SAR11 marine bacteria can be isolated from cryogenically preserved seawater. We reasoned that, since existing SAR11 isolates could be cryopreserved in the presence of 10% glycerol (e.g. 20, 33, 34), then there was no *a priori* reason to believe that natural populations of SAR11 cells could not similarly be preserved. One of many unknown variables, however, was whether or not the process of cryopreservation would result in significant cell loss and thus affect cultivation efficiency. At an equivalent-sized inoculum of five cells, we found that not only were SAR11 strains able to be cultivated from cryopreserved seawater, but the overall culturability was similar between the fresh and cryopreserved seawater samples. Both experiments resulted in the isolation of representatives from the four most abundant SAR11 subclade Ia ASVs in the original inoculum seawater sample, as well as strains from subclade Ib. The cryopreserved seawater sample also proved capable of serving as an inoculum to isolate other SAR11 subclades, as evidenced by the recovery of isolates from within subclade IIIa (two strains) and Va (one strain) from the cryopreserved sample only.

In addition to numerous isolates from SAR11 subclade Ia that appear to represent abundant ASVs in the seawater sample used as inoculum for these experiments, this study yielded seven strains of SAR11 subclade Ib in either mono- or mixed-culture. Despite being a widespread and frequently abundant lineage of SAR11 in the global surface ocean (41–43), only one cultivated representative of subclade Ib had been previously reported, from the Red Sea (28). In addition to the novel isolates of subclade Ib, two strains of subclade IIIa and one of Va were isolated from cryopreserved seawater indicating that a broad range of SAR11 diversity covering at least four major sublineages can be cultivated by this approach, with no apparent negative affect from the cryopreservation treatment itself.

As demonstrated by their recovery here, a range of other oligotrophic marine bacteria can be isolated from cryopreserved seawater coupled with an HTC approach. This includes representatives from the OM60(NOR5) clade, a ubiquitous lineage of oligotrophic marine *Gammaproteobacteria* (OMG) that has been consistently isolated via HTC approaches (e.g. 19, 27, 31), including from coastal Hawai‘i (35). The OM60(NOR5) lineage was the second most-commonly isolated group of marine bacteria, behind only SAR11 subclade Ia, whether using fresh or cryopreserved seawater as inoculum. Of five OM60(NOR5) ASVs present in the seawater used as inoculum, the four most abundant were isolated in both cultivation experiments, indicating no apparent effect of using cryopreserved seawater as an inoculum for isolating members of the OM60(NOR5) lineage. A similar pattern emerged for the *Rhodobacteraceae* lineage of marine *Alphaproteobacteria*, where isolates from the same three ASVs were recovered in each of the two cultivation experiments, out of 13 total *Rhodobacteraceae* ASVs identified in the environmental sample. This included the most abundant *Rhodobacteraceae* ASV from the seawater inoculum, as well as an ASV identical to the previously isolated and genome-sequenced strain HIMB11 from the same sampling location (36). We found only one abundant (> 3) set of strains that was isolated by using raw seawater as inoculum without corresponding strains also isolated using cryopreserved seawater: five strains belonging to two ASVs within the OM43 clade of *Betaproteobacteria* were isolated in mixed- and mono-cultures. While this may indicate that the cryopreservation process had a negative impact on the viability of OM43 clade cells, we note that previously isolated members of this lineage have been successfully cryopreserved in an identical fashion to the method employed in the current study (29, 44). Thus, there is also the potential that this difference stems from stochasticity related to diluting the two inocula nearly one million-fold.

Combining a barcoded next-generation 16S rRNA gene amplicon sequencing approach with a high-throughput dilution culture strategy proved to be a rapid and sensitive means with which to identify strains and assess the constituent taxa within mixed cultures. By barcoding and sequencing each individual culture in the same manner as if it were a mixed microbial community, we obtained taxonomic and proportional data on the microorganisms growing within 58 mixed cultures of up to four constituent ASVs. Recent studies have highlighted the intricacies that interweave the metabolisms of microorganisms inhabiting seawater (e.g. 4, 45); in natural systems, it is probable that a portion of marine microorganisms require as-yet-unidentified growth factors from co-existing cells (46, 47). These dependencies can be identified and investigated by combining a miniaturized, high-throughput approach to cultivate and screen 100s to 1000s of dilution cultures with an inoculum size aimed at growing mixed consortia and a rapid sequence-based screening method that is appropriate for mixed communities, like the one used here.

Consistent with recent observations (27), this set of experiments resulted in the isolation of several bacterioplankton lineages that have been isolated numerous times via HTC and thus appear readily amenable to cultivation via this approach, including members of SAR11 subclade Ia, the OM60(NOR5) lineage, and the *Rhodobacteraceae*. However, it remains that a large portion of the diversity of marine microbes is still being missed in contemporary cultivation efforts. For example, when considering only the putatively heterotrophic, non-cyanobacterial fraction of the microbial community targeted in this study, major lineages including the *Flavobacteriaceae*, SAR86 clade, *Marinimicrobia* (SAR406 clade), and marine Actinobacteria (*Candidatus* Actinomarina) were missed completely. At the single-nucleotide resolution of ASVs, abundant lineages of SAR116 and SAR11 subclades Ib, IIa, and Vb were also conspicuously missed. While this study does not offer a panacea for isolating any of these well-known but as-yet-uncultivated (or undercultivated) lineages in laboratory culture, it presents a method by which high-throughput isolation experiments can be repeatedly performed on an identical set of cryopreserved seawater samples such that requirements for growth can be systematically tested in a cumulative fashion.

In summary, we have demonstrated that a broad range of marine bacterioplankton taxa can be isolated from glycerol-cryopreserved seawater via an HTC approach, and that the cryopreservation process itself did not negatively affect culturability or influence the taxonomic identify of the resulting isolates. Strains of SAR11 subclades Ia, Ib, IIIa, and Va are amenable to isolation from cryopreserved seawater, as well as other abundant lineages of marine bacteria such as OM60(NOR5), oligotrophic *Rhodobacteraceae*, and the PS1 clade. This study demonstrates that cryopreserved seawater can be used as a means to expand the breadth of HTC studies to anywhere cryopreserved stocks can be made, and opens new opportunities to repeatedly interrogate individual water samples or selectively target specific samples for cultivation once ancillary data is in hand.

## Materials and Methods

### Processing of seawater for growth experiments

On 26 July 2017, a 4 L seawater sample was collected in an acid washed polycarbonate (PC) bottle from a depth of 2 m at station STO1 (N 21° 28.974’, W 157° 45.978’) outside of Kāne‘ohe Bay, O‘ahu, Hawai‘i (Fig. 1). Within 1 hr of collection, subsamples of the raw seawater were used to enumerate planktonic microorganisms, cryopreserve subsamples, collect microbial biomass for environmental DNA, and serve as inoculum for a high-throughput cultivation experiment (Fig. 2). Microbial cells were enumerated by staining with SYBR Green I nucleic acid stain (Invitrogen, Carlsbad, CA, USA) and counted on a Guava easyCyte 5HT flow cytometer (Millipore, Burlington, MA, USA) following a previously published protocol (48). To cryopreserve the raw seawater, individual 1.5 mL subsamples were added to 375 µL of 50% glycerol solution (v/v in sterile Kāne‘ohe Bay seawater; 10% final concentration) in 2 mL cryovials (Nalgene, Rochester, NY, USA) at room temperature (24°C), mixed by inverting, and cooled at a rate of −1°C min^−1^ with a Cryo 1°C Freezing Container (Nalgene) inside a −80°C ultracold freezer. Approximately 1.3 L of the raw seawater sample was collected on a 25 mm diameter, 0.1 µm pore-sized polyethersulfone membrane (Supor-100; Pall Gelman Inc., Ann Arbor, MI). The filter was submerged in 500 μL DNA lysis buffer (49, 50) and stored at −80°C until DNA extraction.

### High-throughput cultivation experiment with raw seawater

Growth medium was made following previously published methods (51). Briefly, 20 L seawater samples were collected on 8 July 2017 and subsequently again on 20 September 2017 from a depth of 2 m at station SR4 (N 21° 46.165’, W 157° 78.350’) near Kāne‘ohe Bay, O‘ahu, Hawai‘i (Fig. 1), in acid-washed 4-L PC bottles. Within 1 hr of collection, the seawater was sequentially filtered through pre-rinsed (10 L sterile water followed by 10 L seawater) 0.8-, 0.2-, and 0.1-µm pore-sized polyethersulfone (PES) membranes (AcroPak 20 and Supor 100; Pall Corp., Port Washington, NY, USA) into clean 4-L PC bottles. Bottles were then autoclaved for 3 hours (h) at 121°C and allowed to cool. The sterile seawater was sparged with CO2, followed by air, through three in-line HEPA vent filters (0.3-µm glass-fiber to 0.2-µm PTFE to 0.1-µm PTFE; Whatman, GE Healthcare Life Sciences, Chicago, IL, USA) and stored at 4°C until use. The pH of the seawater was checked prior to autoclaving and after sparging to ensure continuity of the inorganic carbon chemistry.

Subsamples of raw seawater were diluted in the sterile seawater to 2.5 cells mL^−1^ and arrayed in 2 mL volumes (5-cell inoculum) into 576 wells of custom fabricated 96-well Teflon microtiter plates. Plates were sealed with breathable polypropylene microplate adhesive film (VWR, Radnor, PA, USA) and incubated at 27°C in the dark. The presence of cellular growth was monitored at 3.5 and 8 weeks via flow cytometry (Tripp et al. 2008). This experiment is hereafter referred to as HTC2017.

Wells that exhibited positive growth of >10^4^ cells mL^−1^ were sub-cultured by transferring 1 mL into 20 mL of sterile seawater media amended with 400 µM (NH4)2SO4, 400 µM NH4Cl, 50 µM NaH2PO4, 1 µM glycine, 1 µM methionine, 50 µM pyruvate, 800 nM niacin (B3), 425 nM pantothenic acid (B5), 500 nM pyridoxine (B6), 4 nM biotin (B7), 4 nM folic acid (B9), 6 µM myo-inositol, 60 nM 4-aminobenzoic acid, and 6 µM thiamine hydrochloride (B1). Subcultures were subsequently incubated at 27°C in the dark and monitored for growth after 4.5 weeks. Those that again reached >10^−4^ cells mL^−1^ were cryopreserved (500 µL of culture with 10% v/v glycerol, final concentration) in the same manner as described for raw seawater. Cells in the remaining volume of culture (∼18 mL) were collected by filtration through 13 mm diameter, 0.03 µm pore-sized PES membrane filters (Sterlitech, Kent, WA, USA), submerged in 250 µL DNA lysis buffer, and stored at −80°C until DNA extraction.

### High-throughput cultivation experiment with cryopreserved seawater

After 42 weeks of storage at −80°C, one cryopreserved stock of raw seawater from station STO1 was thawed to room temperature (∼24°C), diluted ten-fold in sterile seawater growth medium, and enumerated via staining with SYBR Green I and flow cytometry. The cryopreserved sample was subsequently diluted with nutrient-amended sterile seawater growth medium to two different concentrations: 2.5 and 52.5 cells mL^−1^. The 2.5 cells mL^−1^ dilution was used to create 480 2-mL dilution cultures (5-cell inoculum) in custom fabricated 96-well Teflon microtiter plates, while the 52.5 cells mL^−1^ dilution was used to create 470 2-mL dilution cultures (105-cell inoculum). Ten control wells containing uninoculated sterile seawater growth medium were also included. Teflon plates were sealed with breathable polypropylene microplate adhesive film (VWR) and incubated at 27°C in the dark. Growth was monitored at 2, 3, and 5 weeks post-inoculation as described above. Dilution cultures from the 5-cell inoculum showing positive growth (>10^4^ cells mL^−1^) after 5 weeks of incubation were subcultured by distributing 1 mL of initial culture into 20 mL of sterile seawater growth medium, and monitored for growth during incubation for up to 10 weeks at 27°C in the dark. This experiment is hereafter referred to as HTC2018.

Subcultures from the 5-cell cryopreserved seawater inoculum that reached >10^−4^ cells mL^−1^ were cryopreserved (500 µL of culture with 10% v/v glycerol, final concentration) as described above. Cells in the remaining volume (∼18 mL) were collected by filtration through 0.03 µm pore-sized PES membrane filters (Sterlitech), submerged in 250 µL DNA lysis buffer, and subsequently stored at −80°C until DNA extraction.

The 105-cell inoculum was not subcultured. Instead, wells from one 96-well microtiter plate that exhibited growth (>10^−4^ cells mL^−1^) were cryopreserved by combining glycerol solution to a final concentration of 10% v/v in 250 µL of subculture and frozen as above. Cells in the remaining volume of subculture (∼1 mL) were collected by filtration through 0.03 µm pore-sized PES membrane filters (Sterlitech), submerged in 250 µL DNA lysis buffer, and subsequently stored at −80°C until DNA extraction.

### DNA extraction and sequencing

Genomic DNA was extracted from the environmental sample and all 5-cell subcultures that exhibited growth using the Qiagen DNeasy Blood & Tissue Kit following the manufacturer’s instructions for bacterial cells (Qiagen, Germantown, Maryland, USA). Genomic DNA was extracted from one 96-well microtiter plate of the 105-cell cryopreserved seawater inoculum cultures that exhibited growth using the DNEasy 96 Blood & Tissue Kit (Qiagen) in accordance with the manufacturer’s protocol. Genomic DNA from the environmental sample and all 5-cell subcultures was used as template for polymerase chain reaction (PCR) amplification (Bio Rad C1000 Touch, Bio Rad, Hercules, CA, USA) using barcoded 515F and 926R primers targeting the V4 region of the SSU rRNA gene (52) in a total reaction volume of 25 μL containing 2 μL of genomic DNA template, 0.5 μL each forward and reverse primer, 10 μL 5PRIME HotMasterMix (Quantabio, Beverly, MA, USA), and 12 μL of H2O. The reaction included an initial denaturing step of 3 min at 94°C followed by 40 cycles of 45 sec at 94°C, 1 min at 50°C and 1.5 min at 72°C, and a final extension of 10 min at 72°C.

A nested-PCR approach was used to amplify SSU rRNA gene fragments from genomic DNA recovered from the 105-cell inoculum cryopreserved seawater cultures. The first reaction employed bacterial 27FB (53) and 1492R (54) primers in a 25 μL total reaction volume as described above. The reaction included an initial denaturing step of 3 min at 94°C followed by 35 cycles of 30 sec at 94°C, 1 min at 50°C and 45 sec at 72°C, and a final extension of 18 min at 72°C. PCR products from the first amplification were then used as template for a second amplification reaction using the barcoded 515F and 926R primers (52), using the same reaction conditions as described for the 5-cell inoculum samples.

All PCR products were quantified (Qubit 2.0, Invitrogen), pooled at a concentration of 240 ng sample^−1^, and cleaned (QIAquick PCR Purification Kit, Qiagen). Pooled products were sequenced via three Illumina MiSeq 250 bp paired-end runs using v.2 reagent kits.

### Sequence analysis

The three Illumina MiSeq runs were each separately imported into QIIME2 v2019.4.0, demultiplexed, and paired ends were analyzed for sequence quality and merged (55). The DADA2 software package (56) was then used to denoise sequences, including removal of chimeras. Due to the low quality at the end of the sequences, 10 bases were truncated from the 3’ end of the reverse reads. Sequence reads from the three runs were then merged post-denoising. Amplicon sequence variant (ASV) identities were defined by DADA2 for all reads that varied by at least one base pair. Taxonomy was assigned to each ASV using a Naïve Bayes classifier trained on the Silva rRNA v132 database (57) clustered at 99% similarity and subsequently modified manually based on phylogenetic analyses and the results of previous work. Denoised sequences, ASVs, and taxonomy classifications were imported into R v3.5 (58) using the phyloseq v1.26.1 package (59) for additional manual curation as outlined below. Visualizations were created in R using ggplot2 (60) and in BioVenn (61).

The identities of ASVs found within the cultures and the environmental sample were assigned by QIIME2. For each culture, ASVs represented by fewer than 20 reads were discarded from the data set in order to account for potential sequencing error. Subsequently, the proportion of each ASV in an individual culture was calculated using the read count for that ASV divided by the total reads from the culture, post-curation. Cultures were functionally divided into three separate categories: “monocultures”, “mixed cultures”, and cultures with no discernable, dominant member. All cultures that consisted of ≥90% of reads from a single ASV and contained no other ASVs that were ≥5% of reads were categorized as “monocultures”, and that ASV was assigned a unique isolate identifier in the Hawai‘i Institute of Marine Biology Culture Collection (prefix “HIMB”, followed by unique number). Cultures were defined as “mixed” if they (i) contained an ASV that accounted for <90% but ≥50% of the total reads for that particular culture. This ASV, as well as any other ASV within the mixed culture that contained >5% of the total reads, were assigned unique HIMB identification numbers. (ii) The culture consisted of ≥90% of reads from a single ASV and an additional ASV that was ≥5% of reads. Each of these were also assigned unique HIMB identification numbers. The final category consisted of culture wells that contained no ASVs accounting for ≥50% of the total reads; these were not considered further in the context of this study.

### Analysis of environmental sample

All ASVs represented by <20 reads in the environmental sample were removed in order to account for sequencing error and artifacts. All ASVs that were taxonomically identified as “chloroplast” at the bacterial order-level in the Silva taxonomy were also removed from the data set. The relative abundance of each remaining ASV was calculated as the read count of the individual ASV divided by the total number of reads in the environmental sample, post-curation. Unique identifiers were assigned to all ASVs that remained in the dataset post-curation.

### Culturability statistics

Fundamental culturability statistics were derived as outlined previously (16). Briefly, percent viability (culturability), *V*, is defined as the ratio of the number of viable cells to the total number of cells initially present. It was calculated using the formula:

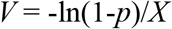

Where *p* is the proportion of wells that scored positive for growth and *X* is the number of cells used for the initial inoculation. To obtain 95% confidence intervals, the exact upper and lower 95% confidence limits for *p* were calculated and inserted back into the original viability equation in place of *p*. The result is the exact upper and lower 95% confidence limits for percent culturability. For this experiment, *p* is defined as the number of cultures that were determined to be either monocultures or mixed cultures as described above. Two separate culturability statistics were calculated: one including mono- and mixed cultures, and one only including monocultures.

### Phylogenetic analyses

Amplicon sequences corresponding to all ASVs were imported into the ARB software package (62) and aligned to a curated database of marine bacterial 16S rRNA gene sequences. Phylogenetic analyses were performed using the RAxML maximum likelihood method with the GTR model of nucleotide substitution under the gamma and invariable-models of rate heterogeneity (63). The heat map of ASV relative abundance was constructed in R v.3.5 (58) using the ggplot2 package (60).

## Data availability

Amplicon sequencing data are available in the Sequencing Read Archive (SRA) under bioproject number xxxxxxxx.

## Acknowledgements

We thank the Center for Genomic Research and Biocomputing at Oregon State University for DNA sequencing, and Dr. Catherine M. Foley for her generous help with the creation of the maps used in Figure 1. This research was supported by funding from the National Science Foundation (grant OCE-1538628 to MSR), the Hawai‘i Institute of Marine Biology (Lord Scholarship fund and graduate research assistantship to EM), and the University of Hawai‘i Marine Biology Graduate Program (to EM). This is SOEST contribution xxx and HIMB contribution xxx.

---

**Figure S1.**
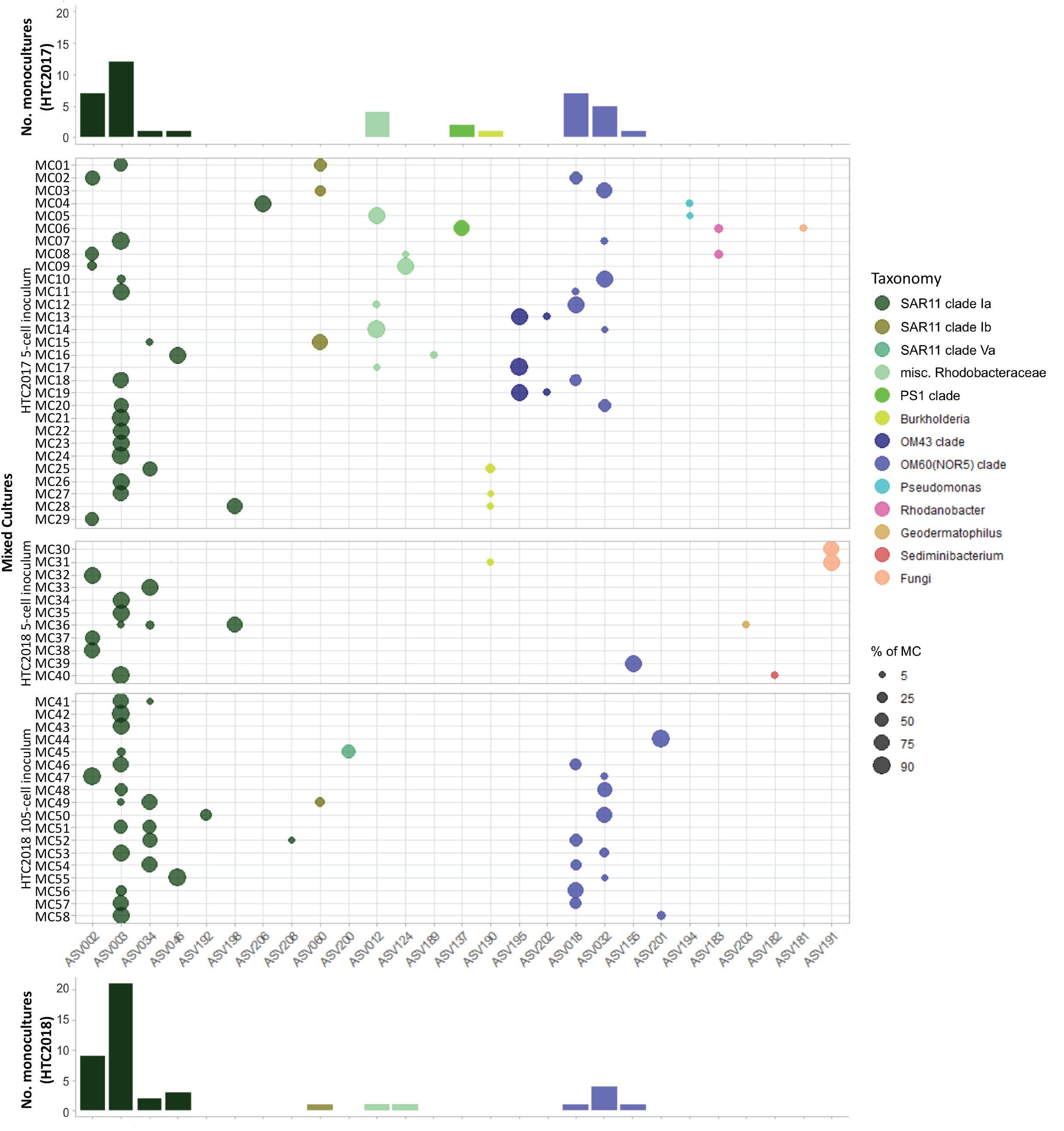
Relative abundance (bubble size) of ASVs identified in mixed cultures from cultivation experiments using fresh seawater (HTC2017, 5-cell inoculum) and cryopreserved seawater (HTC2018 5-cell and 105-cell inocula). Bar charts represent the number of monocultures matching these ASVs cultivated in the two experiments.

## TABLES

**Table S1**. Summary of ASVs, mixed cultures, and isolates recovered in this study.

